# Respiratory bacteria stabilize and promote airborne transmission of influenza A virus

**DOI:** 10.1101/2020.07.23.218131

**Authors:** Hannah M. Rowe, Brandi Livingston, Elisa Margolis, Amy Davis, Victoria A. Meliopoulos, Haley Echlin, Stacey Schultz-Cherry, Jason W Rosch

## Abstract

Influenza A virus (IAV) is a major pathogen of the human respiratory tract where the virus co-exists and interacts with bacterial populations comprising the respiratory microbiome. Synergies between IAV and respiratory bacterial pathogens promote enhanced inflammation and disease burden that exacerbate morbidity and mortality. We demonstrate that direct interactions between IAV and encapsulated bacteria commonly found in the respiratory tract promote environmental stability and infectivity of IAV. Antibiotic-mediated depletion of the respiratory bacterial flora abrogated IAV transmission in ferret models, indicating that these viral-bacterial interactions are operative for airborne transmission of IAV. Restoring IAV airborne transmission in antibiotic treated ferrets by co-infection with *Streptococcus pneumoniae* confirmed a role for specific members of the bacterial respiratory community in promoting IAV transmission. These results implicate a role for the bacterial respiratory flora in promoting airborne transmission of IAV.

## INTRODUCTION

Influenza A viruses (IAVs) are major pathogens of birds and mammals. The primary site of human IAV infection is the upper respiratory tract, with more severe manifestations occurring when the virus accesses the lower respiratory tract. Enhanced IAV morbidity and mortality can also occur due to co-infection with bacterial pathogens, also commonly found in the human upper respiratory microbiota. The best-characterized bacterial synergy of IAV is that with *Streptococcus pneumoniae* (McCullers, 2014; Smith & McCullers, 2014). This synergy operates in multiple aspects of pathogenesis, including enhancing transmissibility of *S. pneumoniae* in murine (A. L. Richard, Siegel, Erikson, & Weiser, 2014) and ferret (McCullers et al., 2010) models.

Recent insights have shown altered pathogenesis resulting from direct bacterial-viral interactions, with the most extensive evidence coming from studies focusing on species found in the gastrointestinal tract. Many classes of enteric viruses, including Picornaviruses (Kuss et al., 2011), Reoviruses (Kuss et al., 2011), and Caliciviruses (Jones et al., 2014), rely on bacteria or bacterial products for infectivity. Such direct interactions have recently been shown to occur also between IAV and respiratory bacteria, including *S. pneumoniae* (David et al., 2019; H. M. Rowe et al., 2019b). These direct interactions enhance the adherence of pneumococcus to cultured respiratory cells *in vitro* and enhanced initial colonization and invasive disease in murine models of otitis and invasive disease (H. M. Rowe et al., 2019b). These interactions also alter host response, as when IAV-pneumococcal complexes were utilized as vaccine antigens, efficacy was greater than that of either vaccine alone (David et al., 2019). While the co-infecting bacterial species typically benefit from IAV infection, the roles for respiratory bacteria on IAV biology remain less well understood. Studies of the respiratory microbiome and susceptibility to IAV infection in household transmission studies have suggested an important role for the composition of the respiratory microbiome in terms of susceptibility to IAV infection (Tsang et al., 2019). Here we demonstrate that IAV directly benefits from interactions with human bacterial respiratory flora, with the bacterial partners conferring environmental stability and enhancing airborne transmission of the virus.

## RESULTS/DISCUSSION

Interactions with bacteria have previously been demonstrated to promote the stability of Picornaviruses (Aguilera, Nguyen, Sasaki, & Pfeiffer, 2019; Robinson, Jesudhasan, & Pfeiffer, 2014) and Reoviruses (Berger, Yi, Kearns, & Mainou, 2017; Robinson et al., 2014). We hypothesized that bacterial stabilization of environmental IAV via direct interactions may be one mechanism the virus exploits to retain infectivity following release into the environment. To test this, bacterial cultures from several respiratory tract–colonizing pathogens, previously shown to associate with influenza A viruses (David et al., 2019; H. M. Rowe et al., 2019b), were incubated with influenza virus strain A/Puerto Rico/8/1934(H1N1) (PR8), centrifuged, and washed to remove nonadherent virus. The co-sedimented material was subjected to desiccation in a speed vac and then rehydrated to determine viral infectivity. Speed vac mediated desiccation, while not biologically relevant, allows concentration of bacterial and viral particles and ability to measure log fold changes in viral viability promoted by the bacterial complex. Real world conditions would subject the bacterial-viral complex to less harsh desiccation stressors and furthermore would be in the context of host derived molecules.

Virus desiccated in the presence of *S. pneumoniae* (pneumococcus) or *Moraxella catarrhalis* retained viability and infectivity, whereas IAV complexed to *Staphylococcus aureus, Staphylococcus epidermidis*, non-typeable *Haemophilus influenzae*, or *Pseudomonas aeruginosa* did not retain infectivity of IAV (Figure 1A). The desiccation resistance conferred by the IAV-bacterial complex was independent of bacterial viability, as ethanol-killed pneumococci, or *ΔspxB* mutant, which maintains higher desiccation viability than wild-type pneumococcus (H. M. Rowe et al., 2019a), promoted influenza viability to an equivalent degree as live wild-type *S. pneumoniae* (Figure 1B). However, pneumococcus had to be intact to promote infectivity of IAV, as virus co-sedimented in the presence of pneumococci killed and lysed with β-lactam antibiotics retained significantly less viability than virus desiccated in the presence of live pneumococci (Figure 1B). These data indicate that direct interactions between IAV and respiratory bacteria can promote environmental stability of IAV during desiccation in a species-specific manner and that bacterial association alone is not sufficient to stabilize IAV.

**Figure 1:**
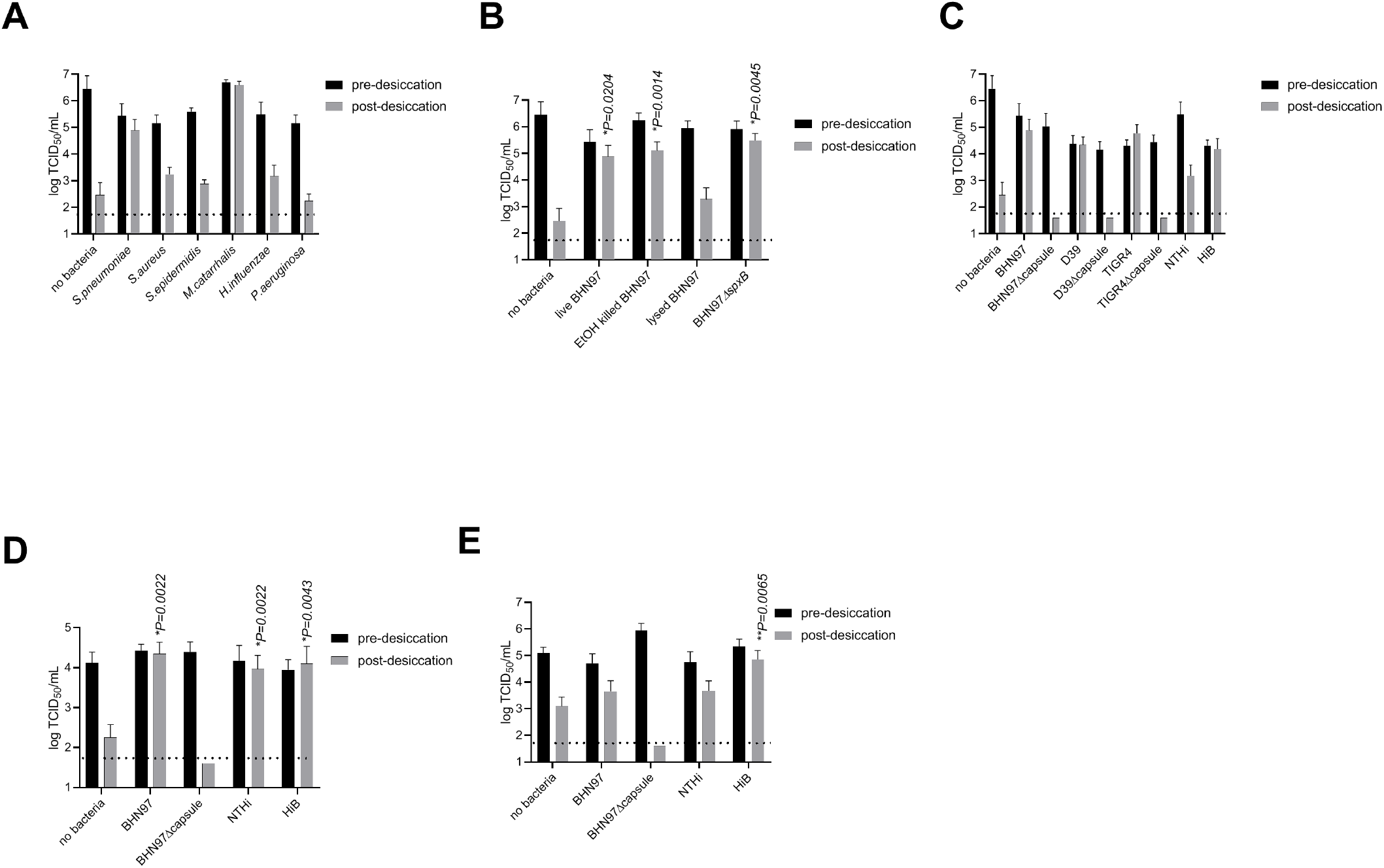
Nasopharyngeal bacteria promote influenza virus desiccation stability. Indicated bacterial strain was preincubated with influenza virus PR8 (A-C) or A/California/04/2009(H1N1) (D), or A/Wisconsin/67/2005 (H3N2) € followed by centrifugation and washes to remove non-associated virus and desiccation in a speed vac (grey) or not subject to desiccation (black). Virus alone was in a small volume (less than 10 μL) of either cell culture media or egg fluid and was directly dessicated in the speed vac. (A) Nasopharyngeal tract–colonizing bacteria provide differing degrees of IAV desiccation protection. (B) Pneumococcal viability does not affect desiccation promotion of IAV, as ethanol-killed or *ΔspxB*, pneumococcal mutant with enhanced desiccation tolerance, had equivalent protection of IAV infectivity as live pneumococci. β-lactam–killed and lysed pneumococci did not promote viability retention. (C) Desiccation survival of IAV with encapsulated and non-capsulated strains of *S. pneumoniae* and *H. influenzae*. (D) Pneumococcal capsule and *H. influenzae* promote stability of A/California/04/2009 (H1N1). (E) *H. influenzae* serogroup B promotes stability of A/Wisconsin/67/2005(H3N2). Bars represent mean and error bars represent standard deviation of at least 6 biological replicates. *P* values calculated by Mann-Whitney testing compared to virus desiccated in the absence of bacteria; dotted line represents limit of detection.

The extensively hydrated polysaccharide capsule can promote environmental survival of *S. pneumoniae* (Hamaguchi, Zafar, Cammer, & Weiser, 2018). Further, lyophilization of live-attenuated influenza vaccine in sugar-containing solutions promotes maintenance of viability (Lovalenti et al., 2016). We hypothesized that the polysaccharide capsule may be one mechanism by which direct interactions between IAV and the pneumococcus promotes viral stability. Targeted deletions of the capsule locus were made in three pneumococcal strains, representing distinct three genotypes and serotypes. These mutants, and parental strains, were incubated with IAV strain PR8, co-sedimented and subjected to desiccation. Capsule made no discernable impact on the initial association and adherence of IAV to the bacterial cells, similar to previous studies (H. M. Rowe et al., 2019b), but only virus co-sedimented in the presence of encapsulated pneumococcal strains retained infectivity. This phenomenon was not specific to pneumococcal capsule, as IAV co-sedimented in the presence of encapsulated *H. influenzae* serotype B strain (HiB) demonstrated significantly enhanced viability compared to that of IAV co-sedimented with non-typeable *H. influenzae* (Figure 1C). These data indicate an important role for the polysaccharide capsule of *S. pneumoniae* and *H. influenzae* in conferring desiccation tolerance to IAV.

To confirm that findings were also operative in human relevant pathogens, desiccation experiments were performed with both A/California/04/2009 (H1N1) and A/Wisconsin/67/2005 (H3N2) desiccated in the presence of capsular and non-encapsulated respiratory bacteria. Similar to PR8, co-sedimentation of A/California/04/2009 with *S. pneumoniae* resulted in significantly enhanced desiccation tolerance, a phenotype that was dependent upon expression of the pneumococcal polysaccharide capsule (Figure 1D). However, both the non-typeable *H. influenzae* and HiB promoted desiccation tolerance of A/California/04/2009, suggesting that the role of *H. influenzae* serotype B capsule is less important in promoting desiccation tolerance of A/California/04/2009 compared to PR8 and that another *H. influenzae* surface factor may play a role in stabilizing A/California/04/2009. Interestingly, the desiccation tolerance of A/Wisconsin/67/2005 (Figure 1E) was not promoted by *S. pneumoniae* regardless of capsule status, but was promoted by encapsulated *H. influenzae* serotype B. Taken together, these data suggest that human respiratory bacteria are capable of stabilizing H1N1 strains of IAV but this stabilization may be subtype specific, with certain respiratory bacteria stabilizing certain IAV subtypes.

These observations of respiratory tract–colonizing bacteria conferring desiccation tolerance to IAV suggests a mechanism whereby, during shedding from an infected host, the viral particles associated with specific members of the respiratory microbial community may have enhanced environmental stability and, hence, transmissibility. First, to determine if we could alter the respiratory microbial community, ferret anterior nasal swabs were collected, a subset of these animals were treated immediately after sample collection and three days later by application of mupirocin ointment, commonly used in nasal decolonization prior to surgery (Septimus, 2019), to the ferret nostrils using a polyester tipped swab to apply the ointment to the exterior of the nares and interior of the nostrils to the depth of the first turbinate. All ferrets were sampled again 24 hours after the final treatment to determine the impact of mupirocin on respiratory bacterial communities. DNA was extracted from swabs and the V3-V4 region of 16S was sequenced to determine microbial population composition. While total bacterial burden was significantly reduced as determined by 16S rRNA copies per swab (Figure 2A), the overall community diversity was not significantly altered (Supplementary Figure S1). The decrease in bacterial burden was limited to particular species, including multiple gram positive cocci and *Moraxella*, which were significantly reduced in treated but not control animals (Figure 2B and Supplemental Figure 2).These data indicate that mupirocin treatment selectively reduced the relative burden of multiple respiratory bacterial species, including *Streptococcus* and *Moraxella*, both of which mediate IAV binding and desiccation tolerance.

**Figure 2:**
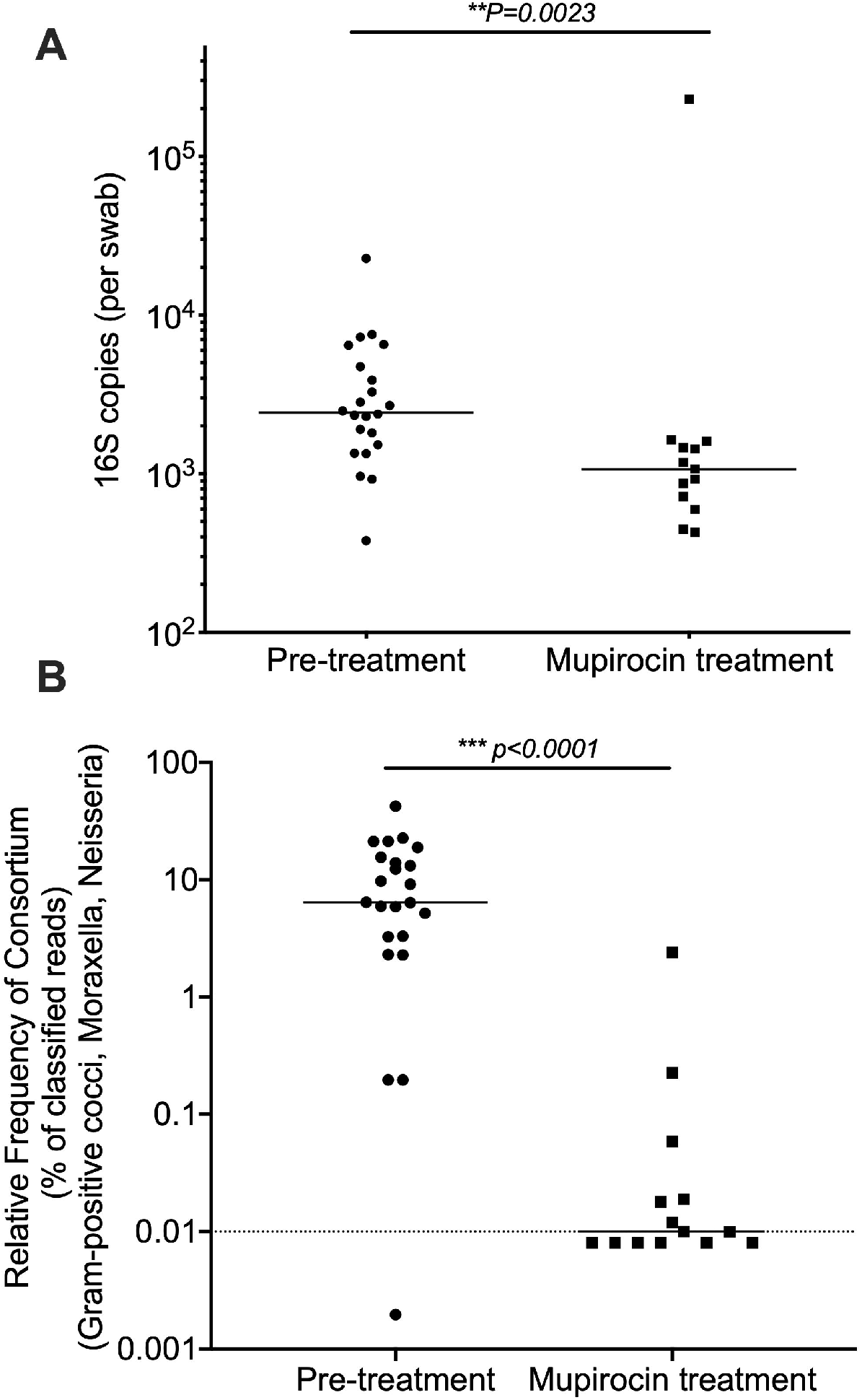
Impact of mupirocin on ferret respiratory microbial community composition. (A) Bacterial content of the nasal passages is significantly lower after mupirocin treatment as measured by bacterial 16S copies recovered on nasal swabs prior to and following treatment. (B) Microbiome content of Gram-positive and Gram-negative cocci (specifically the relative frequency of Moraxella, Neisseria, Lactococcus, Vagococcus, Enterococcus hirae, Streptococcus fryi and suis) is significantly reduced following treatment. Each dot represents data from a swab collected from an individual ferret. Solid line indicates median for each group, dashed line represents limit of detection. Groups compared by Mann-Whitney test.

Based on the observations that specific bacterial species can mediate IAV infectivity during desiccation and mupirocin depletes bacterial species from the nasal passages, we hypothesized that mupirocin treatment would adversely impact IAV transmission. To determine the effect of this community disruption on airborne transmission of IAV, ferrets were treated by application of mupirocin ointment to the ferret nostrils, on days one and three prior to viral infection and at each nasal wash collection point, with ointment administered after collection of nasal wash. Pairs of donor and aerosol-contact ferrets were housed in cages with perforated dividers such that the ferrets could not directly contact each other. Donor ferrets were infected with A/California/4/2009 (H1N1) by the intranasal route. Nasal wash samples were collected on days 3, 5, 7, 9 and 11 or 12 post viral challenge from both donor and contact animals to monitor viral burden. Control ferrets with no manipulation of the respiratory microbial community had a 75% transmission rate, with two of four ferrets having culturable virus in their nasal washes and an additional contact becoming seropositive by day 21 post challenge (Figure 3A,B). Depleting the respiratory microbial community by applying mupirocin to the nostrils of donor and contact ferrets completely abrogated airborne transmission, with no contact animals expressing positive viral titers nor seroconverting (Figure 3C,D). Treatment of only the donors was sufficient to block airborne viral transmission with no contact animals with positive viral titers or seroconversion (Figure 3E,F). All directly infected animals regardless of treatment status shed similar viral loads, seroconverted to infection, and exhibited similar clinical symptoms (Supplementary Table 1). These data indicate that perturbation of bacterial communities in the respiratory tract of donor ferrets by mupirocin treatment results in reduced airborne transmission of A/California/4/2009 (H1N1) influenza virus.

**Figure 3:**
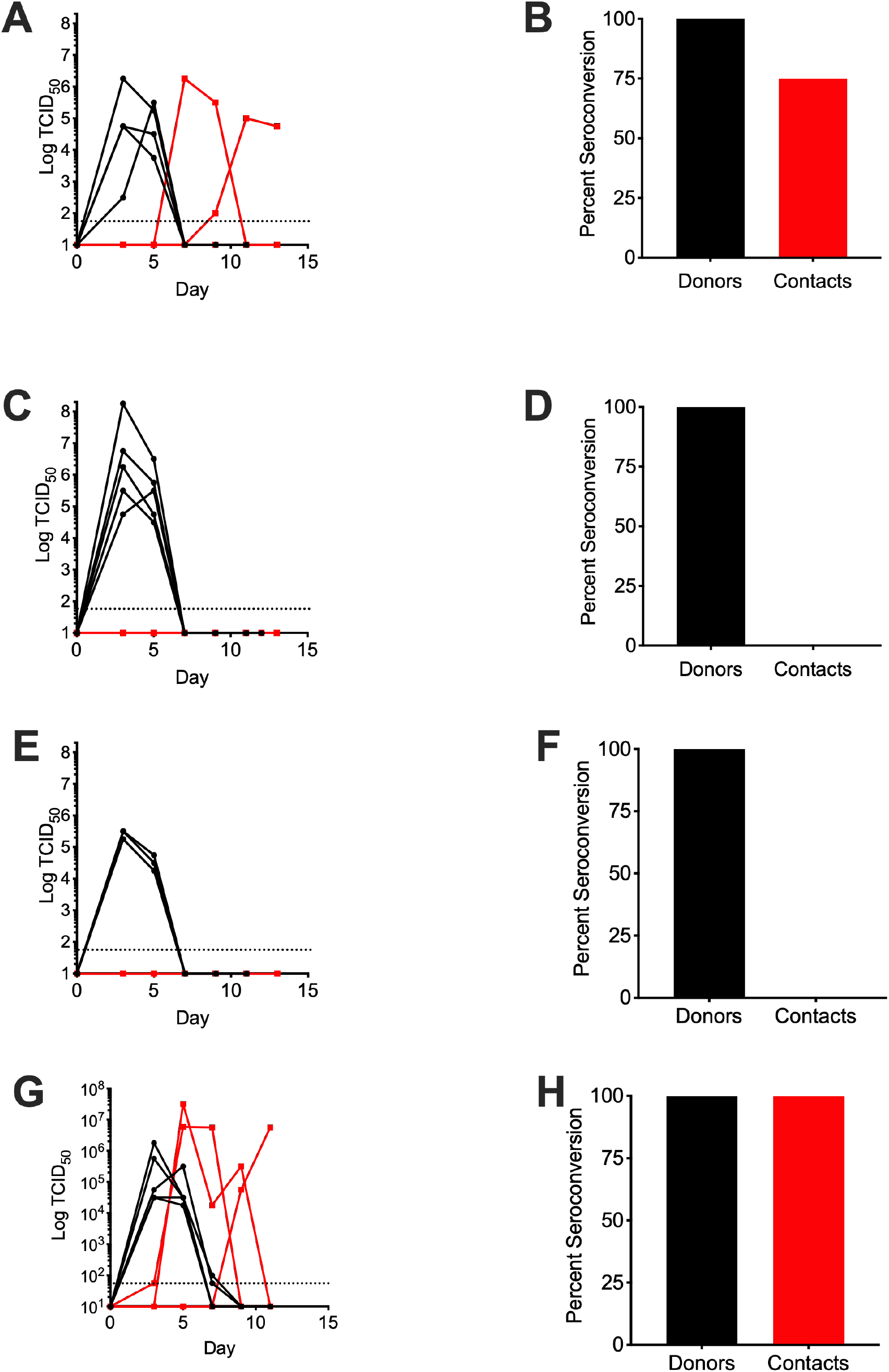
Nasopharyngeal bacteria promote airborne transmission of influenza virus. Donor ferrets were infected with influenza A virus A/California/04/2009 (H1N1) and paired 24 hours post infection with aerosol-contact ferrets in the same cage with perforated dividers separating the animals. (A,C,E,G) Influenza virus burden in nasal lavage measured by 50% tissue culture infectious dose (TCID_50_). (B,D,F,H) Percent of animals who seroconverted as measured by hemagglutination inhibition (HAI) assay titer greater than 1:80 dilution by day 21 post infection. Donors=black, aerosol contacts=red. Dotted line represents limit of detection for TCID_50_ assay, Days= days post infection of donor animals. (A,B) Ferrets with no manipulation of the respiratory microbiota, n=4 donors and 4 contacts. (C,D) Both donor and contact ferret nostrils treated with mupirocin ointment, n=5 donors and 5 contacts. (E,F) Donor ferret nostrils treated with mupirocin ointment, n=3 donors and 3 contacts. (G,H) Donor ferret nostrils treated with mupirocin followed by colonization with 10^6^ CFU of *S. pneumoniae* strain BHN97Mup^R^Strep^R^. Each data point represents an individual ferret over time.

If our hypothesis that respiratory bacteria promote IAV transmission is correct, then restoring bacterial communities that both bind to and stabilize IAV should rescue IAV transmission in the mupirocin-treated donor animals. Because *S. pneumoniae* was shown to stabilize IAV from desiccation stress and effectively colonizes the respiratory tract of ferrets, a separate group of mupirocin-treated donor ferrets was colonized with mupirocin-resistant *S. pneumoniae* 2 days after IAV challenge, when viral shedding is at near peak levels (Roberts, Shelton, Stilwell, & Barclay, 2012). Mupirocin treatments were identical to those undertaken in Figure 3. Pneumococcal colonization was robust and stable throughout viral sample collection across donor animals (Supplemental figure 3). Viral transmission was restored upon colonization of *S. pneumoniae*, with 60% of contact animals having viable virus in their nasal washes and all contact ferrets seroconverting to infection (Figure 3G,H). While donor viral loads were similar to previous groups, donor symptoms were higher in colonized ferrets (Supplementary Table 1), so we cannot rule out an additional effect from enhanced donor symptoms on shedding to contacts, However, these data suggest that colonization of donor ferrets by *S. pneumoniae* is sufficient to rescue the IAV transmission defect resulting from mupirocin depletion of the respiratory flora.

Viral-bacterial synergies are inherently complex with interactions between both bacteria, virus, and the host immune system being operative in synergistic as well as antagonistic relationships. We demonstrated that common members of the human nasopharyngeal microbiome including *S. pneumoniae, M. catarrhalis*, and *H. influenzae* can enhance desiccation survival of H1N1 influenza A viruses when virus was in complex with the bacterial surface. In our study desiccation survival was enhanced by the presence of bacterial capsule in both *S. pneumoniae* and *H. influenzae* suggesting the polysaccharide capsule plays an important role in retaining IAV infectivity under these conditions. These findings reflect studies of enteric viruses such as Picornaviruses whose stability can be enhanced via direct bacterial interactions(Aguilera et al., 2019) or interaction with bacterial lipopolysaccharide (Robinson et al., 2014). Bacterial lipopolysaccharide and peptidoglycan also enhance thermostability of Reoviruses, which in turn promotes infection of host cells following environmental stress (Berger et al., 2017). Similar virion stability and infectivity enhancement has also been observed with human Astroviruses (Perez-Rodriguez et al., 2019). This suggests that interactions between viral pathogens and the bacterial communities are likely operative at distinct host niches including the respiratory tract in addition to the better characterized synergies operative for enteric pathogens.

The environmental persistence of IAV is dependent upon multiple factors and can vary considerably between viral subtypes (Kormuth et al., 2019). The differing capacity of respiratory bacterial species to promote stabilization of the H1N1 viruses versus the H3N2 subtype suggests that there may be additional important differences between distinct IAV subtypes for their capacity to be stabilized by direct bacterial interactions with some IAV subtypes requiring distinct bacterial species for binding and stabilization. Further, some IAV strains may not be stabilized by human respiratory bacteria, but instead by bacteria found in the natural reservoir of the respective IAV strain. Additionally, the respiratory microflora of model organisms may impact IAV transmissibility, underscoring the potential importance of the native bacterial flora when investigating IAV transmissibility. It should also be noted that we utilized relatively young ferrets aged 8-10 weeks, while many other investigations querying influenza transmission that routinely utilize ferrets of 4-12 months of age (Belser, Eckert, Tumpey, & Maines, 2016; M. Richard et al., 2020; T. Rowe et al., 2010). The rationale for the utilization of younger ferrets was primarily due to previous work demonstrating ferrets in this age group rapidly transmit *S. pneumoniae* by both contact and aerosol routes (McCullers et al., 2010; H. M. Rowe et al., 2019a). Whether our findings would extend to older ferrets that likely have distinct microbial community composition remains an unknown but intriguing question. The relevance of results to various IAV challenge doses is also an important question, as bacterial-mediated synergies may only be important at specific viral thresholds of infectivity.

These findings suggests that, unlike the enteric microflora, which enhance viral infectivity of the same host, the respiratory microflora of the infected host is primarily operative in viral infectivity of the subsequent host. In household transmission studies of IAV, *S. pneumoniae* or closely related streptococcal species were identified in approximately 95% of samples collected both the child index cases and the household contacts who developed influenza (Zhang et al., 2020). These data suggest that modulating donor respiratory flora via antibiotic exposure or vaccination may profoundly affect IAV transmission. It should be stressed that in our study topical antibiotics were given prior to IAV challenge with no impact on disease severity in the donor animals. Even in the light of this limitation, targeting bacterial-mediated transmission may represent a novel strategy of IAV infection control that could be explored.

## MATERIALS AND METHODS

### Ethics Statement

All experiments involving animals were performed with approval of and in accordance with guidelines of the St. Jude Animal Care and Use Committee. The St. Jude laboratory animal facilities have been fully accredited by the American Association for Accreditation of Laboratory Animal Care. Laboratory animals were maintained in accordance with the applicable portions of the Animal Welfare Act and the guidelines prescribed in the DHHS publication *Guide for the Care and Use of Laboratory Animals*.

### Bacterial and viral strains and growth conditions

The *S. pneumoniae* strains BHN97 (serotype 19F), D39 (serotype 2), and TIGR4 (serotype 4) were inoculated onto tryptic soy agar (TSA) (GranuCult-Millipore Burlingon MA) plates supplemented with 3% sheep blood (iTek StPaul MN) and 20 μg/mL neomycin (Sigma St Louis MO) and then grown overnight at 37°C in a 5% CO_2_ humidified incubator. Strains were then inoculated directly into Todd-Hewitt (BD Franklin Lakes NJ) broth supplemented with 0.2% yeast extract (BD) (ThyB) and grown to log phase for use in experiments. A capsule mutant was generated by transforming SPNY001 genomic DNA containing a Sweet Janus cassette that replaces the capsule locus (Grijalva et al., 2014) into strains BHN97, D39, and TIGR4 and confirmed by the lack of latex bead agglutination (Statens Serum Institute Copenhagen Denmark) for the respective capsule.

The nontypeable *H. influenzae* 86-028NP(Harrison et al., 2005), originally isolated from a patient with chronic otitis media, and the encapsulated *H. influenzae* serotype b strain 10 211(ATCC) were grown on chocolate agar supplemented with 11,000 units/L bacitracin (BD) and then directly inoculated into brain heart infusion broth (BD) supplemented with 0.2% yeast extract (BD), 10 μg/mL hemin, and 10 μg/mL NAD and grown with aeration to mid-log phase. *Staphylococcus aureus* strain USA400, *Staphylococcus epidermidis* strain M23864:W2 (ATCC), *P. aeruginosa* Xen41 (PerkinElmer), and *Moraxella catarrhalis* (Helminen et al., 1994) were grown on unsupplemented TSA plates, directly inoculated in brain heart infusion broth supplemented with 0.2% yeast extract, and grown with aeration to mid-log phase for use in experiments. The influenza A virus A/Puerto Rico/8/1934 (PR8) and A/Wisconsin/67/2005 (H3N2) were grown in Madin-Darby canine kidney (MDCK) cells. The A/California/4/2009 virus was grown in allantoic fluid of 10- to 11-day-old embryonated chicken eggs. PR8 is of unknown passage history, A/Wisconsin/67/2005 is a 2^nd^ cell passage from a third egg passage, A/California/4/2009 is a 5^th^ egg passage.

For ferret pneumococcal colonization, *S. pneumoniae* strain BHN97 was made mupirocin- and streptomycin-resistant (BHN97 Mup^R^Strep^R^) to enable continued treatment of ferrets with mupirocin ointment and collection of nasal lavage with streptomycin to reduce risk of aspiration pneumonia during ketamine sedation and nasal wash collection. Streptomycin-resistance was conferred via mutation of *rpsL* (TIGR4 Sp_0271) by introduction of a K56T mutation(Martin-Galiano & de la Campa, 2003) generated by splicing overlap extension (SOE) PCR using two fragments that each had the point mutation. The first PCR fragment amplified 969 bp upstream and the first 180 bp of *rpsL* using primers RpsL_Up_F (GCCGTAGTCATCTTTCTTGGCATC)/ RpsL_Up_R(CTGAGTTAGGTTTTGTAGGTGTCATTGTTC). The second PCR fragment amplified bp 151 to 414 of rpsL plus 752 bp downstream using primers RpsL_Down_F(GAACAATGACACCTACAAAACCTAACTCAG)/ RpsL_Down_R(CTAATTTGAACCCGGGCTAAAGTTAG). The entire SOE PCR product was amplified using RpsL_Up_F/ RpsL_Down_R and was transformed into strain BHN97; resistant mutants were selected for on TSA supplemented with 3% sheep blood and 800 μg/mL streptomycin. Mupirocin-resistance was spontaneously generated and selected for by plating turbid culture of BHN97 *rpsL_K56T_* on TSA supplemented with 3% sheep blood, 800 μg/mL streptomycin, and 10 μg/mL mupirocin and then selecting spontaneously resistant colonies.

### Co-sedimentation and desiccation

Co-sedimentation was performed as previously described (H. M. Rowe et al., 2019b). Briefly, mid-log bacterial cultures were washed and normalized to 10^8^ CFU/mL in phosphate-buffered saline (PBS). 3×10^7^ TCID_50_ (50% tissue culture infectious dose) influenza virus was added and samples rotated 30 minutes at 37°C. Samples were centrifuged and washed twice with PBS. Samples not subjected to desiccation were immediately resuspended in 100 μL 1x penicillin/streptomycin solution (Gibco) and frozen at −80°C for viral quantification. Samples designated for desiccation were spun for 60 minutes in a Speed Vac until the pellet was dry. Pellets were resuspended in 100 μL 1x penicillin/streptomycin solution and frozen at −80°C for viral quantification. Viral titers were determined by TCID_50_ on MDCK cells (Cline et al., 2011). Three to six biological replicates were performed for each strain.

Ethanol-fixed pneumococci were prepared by resuspending 10^8^ CFU BHN97 in 1 mL of ice-cold 70% ethanol for 5 minutes on ice. Cells were pelleted, supernatant was removed, and pellets were dried at 55°C for 5 minutes to remove residual ethanol. Viability loos was confirmed by plating on TSA/blood. β-lactam–killed pneumococci were prepared by resuspending 10^8^ CFU BHN97 in 1 mL 10x penicillin-streptomycin (Gibco) solution in PBS and incubating 30 minutes at 37°C. Viability loss was confirmed by plating on TSA/blood, and lysis was confirmed by microscopy examination.

### Ferret infection

All ferrets were maintained in BSL2, specific pathogen–free facilities. Microbiome collection swabs, treatment of nostrils with ointment, infection and blood collections were conducted under general anesthesia with inhaled isofluorane at 4%. Nasal washes were collected under ketamine sedation following intramuscular injection of ketamine to the thighs of restrained ferrets. All ferrets were monitored twice daily for symptoms during infection. Weights were measured daily and temperature collected daily from implanted microchips.

Nine-week-old male castrated ferrets (Triple F Farms Gillet PA), confirmed to be seronegative for Influenza A viruses (seronegativity to Influenza B viruses not tested) prior to start of study, were housed two per cage, separated by a perforated barrier. Experimental groups had 3 to 5 donor-contact pairs. Animals designated for treatment with mupirocin ointment had 75mg 2% mupirocin in polyethylene glycol (Perrigo) applied to exterior of nostrils and interior of anterior nares up to the first turbinate with a polyester applicator swab (Puritan) three days prior to, one day prior to, on the day of infection (post-instillation of virus inoculum), and on each sampling day (post collection of nasal lavage); untreated animals were not treated at those time points. Donor animals were infected with 10^6^ 50% tissue culture infectious doses (TCID_50_) of influenza A virus A/California/04/2009 (H1N1) in 1 mL of phosphate-buffered saline (PBS), instilled equally between both nostrils. Contact animals were introduced the cages, separated with a perforated divider, 24 hours after infection of the donors. On days 3, 5, 7, 9, and 11 or 12 post infection of donor animals, donor and contact ferrets were sedated with ketamine and nasal lavage was collected in 1 mL PBS supplemented with 1x penicillin/streptomycin (Gibco) divided equally between each nostril. Nasal lavage was stored at −80°C for viral quantification. Animals designated for co-infection with *S. pneumoniae* were treated with mupirocin ointment and infected as described above. Then on day 2 post influenza infection, *S. pneumoniae* strain BHN97 Mup^R^Strep^R^, grown as described in the co-sedimentation and desiccation section, was normalized to 5×10^6^ CFU per 600 μL PBS and instilled equally between both nostrils. Samples were collected as above, except penicillin was omitted from PBS. Prior to storage at −80°C, 30μL of nasal wash was removed and serially diluted in PBS and plated on TSA supplemented with 20 μg/mL neomycin and 3% sheep blood for bacterial quantification. Viral titers were determined by TCID_50_ on MDCK cells (Cline et al., 2011). Briefly, MDCK cells were infected with 100 μL 10-fold serial dilutions of sample and incubated at 37°C for 72 hours. Following incubation, viral titers were determined by hemagglutination assay (HA) using 0.5% turkey red blood cells and analyzed by the method of Reed and Munch (Reed & Meunch, 1938). For samples that were negative by HA, residual supernatant was removed and wells were washed once with PBS and then stained for one hour at room temperature with 0.5% crystal violet in a 4% ethanol solution. Wells were washed with tap water and infected wells determined by destruction of the monolayer. TCID_50_ was again determined using the method of Reed and Munch as above. On days 14 and 21 post IAV challenge, ferrets were sedated with isoflurane, and 1 mL blood was drawn from the jugular vein. Blood was allowed to clot overnight at 4°C. Serum was collected following centrifugation to pellet clot and stored at −80°C. For determination of seroconversion, serum was treated with RDE (Hardy Diagnostics) overnight at 37°C. RDE was inactivated via incubation at 56°C for 1 hour, followed by dilution in PBS for a final dilution of 1:4, and freezing at −80°C for at least 4 hours to continue to inactivate neuraminidase. Starting with a 1:40 dilution of sera and serial 2-fold dilutions in PBS, sera were mixed with 4 hemagglutination units of A/California/04/2009 and incubated 30 minutes at room temperature. Following incubation, an equal volume of 0.5% washed turkey red blood cells in PBS was added to each well and incubated a further 60 minutes at 4°C. Hemagglutination inhibition titer was read as the most dilute well with a negative hemagglutination reaction.

### Microbiome analysis

Prior to treatment on day −3 and again on day of infection, just prior to the infection, the microbiome was sampled from the anterior nares of all ferrets. A flocked polyester swab (Copan, flexible minitip) was inserted into the ferret nares to the depth of the first turbinate and the interior of each nostril was swabbed for 15 seconds per nostril, before insertion of the same swab into the other nostril. Swabs were stored dry at −80°C for DNA preparation. DNA was extracted from nasal swabs after resuspension using methods to improve the bacterial species captured in low abundance samples (Davis et al., 2019). Bacterial DNA content was assessed using a 16S rRNA quantitative PCR with a plasmid containing E. coli 16S gene as the standard, using previously described primers and probe (Nadkarni, Martin, Jacques, & Hunter, 2002) and Fast Universal PCR Master Mix (TaqMan) supplemented with 3 mM MgSO_4_. Microbiome 16S rRNA gene amplification was performed using ‘touch-down’ PCR cycling of V3-V4 amplicon (Dickson et al., 2014) and sequencing was performed as previously described (Golob et al., 2017) at the St. Jude Hartwell Center. Classification of reads was done based on phylogenetic placement on reference tree; briefly: Illumina MiSeq paired-end reads were run through DADA2 pipeline (Callahan et al., 2016) (version 1.10.1) to correct sequencing errors and determine amplicon sequence variants (ASVs). These ASVs were then used to recruit full-length 16S rRNA gene sequences from Ribosomal Database Project release 16.0(Cole et al., 2014) to construct a phylogenetic reference dataset and tree. The amplicon sequences were then placed onto the reference tree using pplacer (Matsen, Kodner, & Armbrust, 2010). Sequence reads are available through NCBI through accession # (to be provided upon publication).

### Statistical analysis

All tests were performed with GraphPad Prism7. Comparisons were made via Mann-Whitney testing, with a *P* value of less than 0.05 considered significant.

## ACKNOWLEDGEMENTS

JWR is supported by 1U01AI124302 and 1RO1AI110618, SSC by NIAID contract HHSN272201400006C, and all by ALSAC. The content is solely the responsibility of the authors and does not necessarily represent the official views of the National Institutes of Health.

## Author contributions

HMR, BL, VAM, AD and HE performed the experiments. HMR, SSC, and JWR designed the study. HMR, EM and JWR analyzed the data. HMR and JWR wrote the manuscript, and all authors edited and approved the final manuscript.

## Materials & Correspondence

All materials and data will be made available upon request to.

