## Supplemental Data for "Respiratory bacteria stabilize and promote airborne transmission of influenza A virus"

SUPPLEMENTAL FIGURES

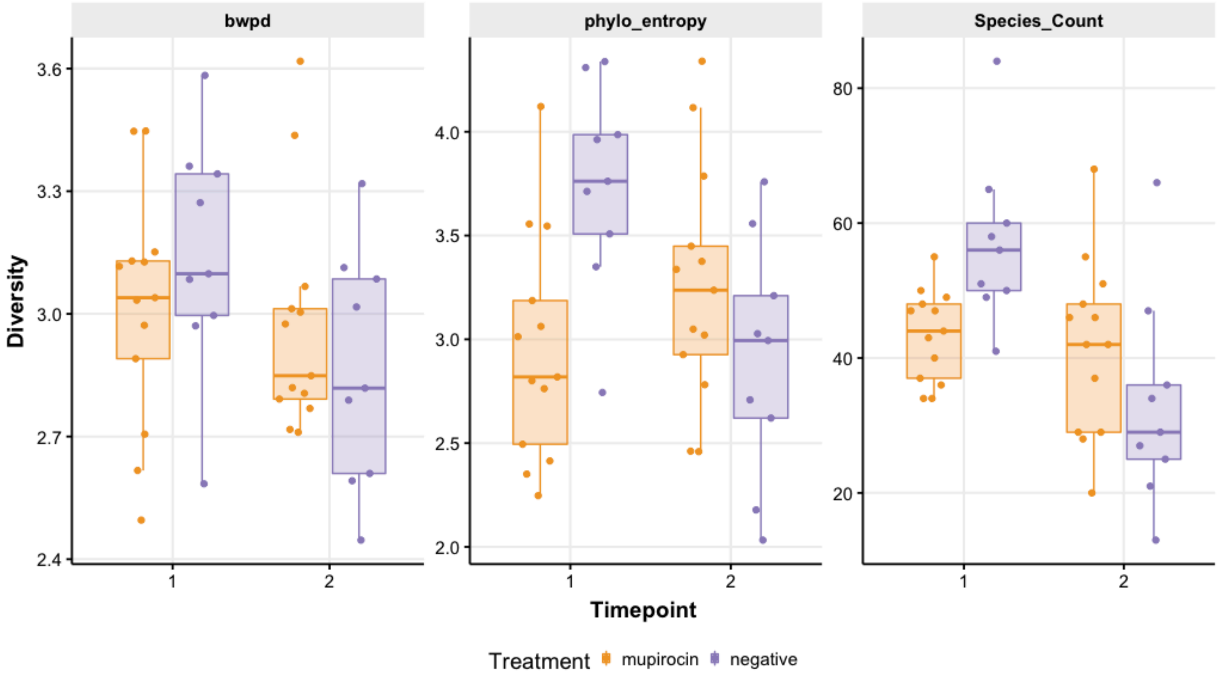


Figure S1: Nasal microbiome diversity after mupirocin treatment. DNA extracted from nasal swabs were examined by 16S rRNA amplicon sequencing of V3-V4. Microbial diversity as measured by the 3 methods indicated at the top of each graph shows that the indices' levels do not significantly change after mupirocin treatment.

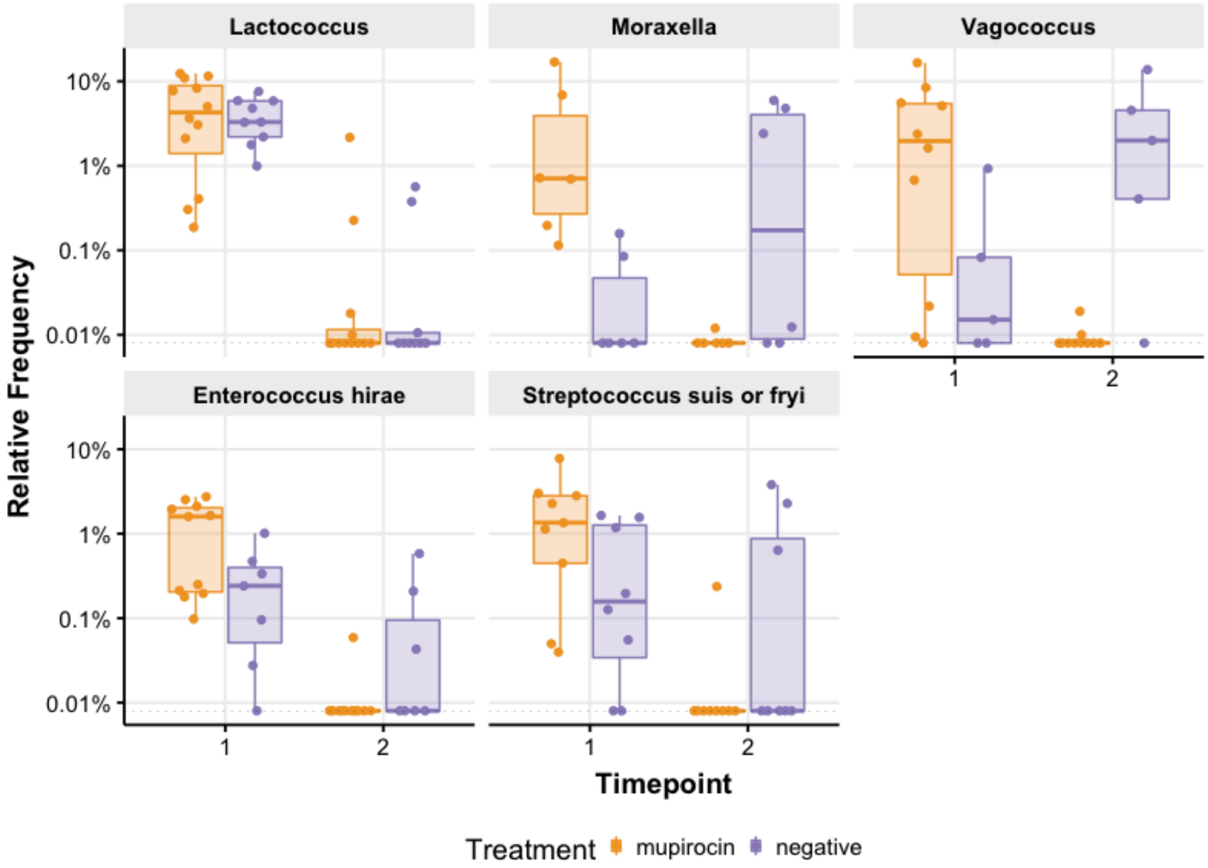


Figure S2: Alterations of mupirocin sensitive microbes in nasal microbiome composition after mupirocin treatment. Gram positive cocci (*Lactococcus, Vagococcus*, *Enterococcus hirae, Streptococcus fryi* and *suis)* and *Moraxella* were commonly eradicated to below the limit of detection (dotted line) after treatment with mupirocin.

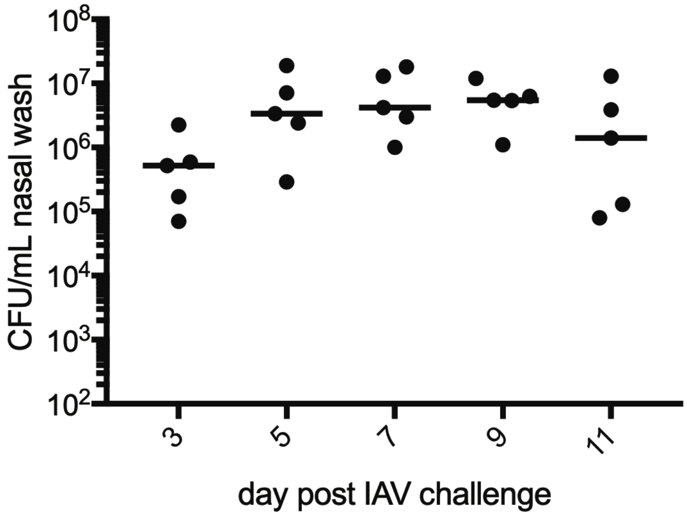


Figure S3: *S. pneumoniae* colonization of donor ferrets remains stable over infection and mupirocin treatment. Each dot is one animal, line is median.

Supplemental Table 1: Symptom scores of donor ferrets

| Ferret ID | Max Weight loss (% starting weight) | Max temperature, fever[a] bold text | Max lethargy score[b] | Days with Sneezing | Days with nasal/ocular discharge |
|---|---|---|---|---|---|
| Untreated 1 | 5.8 | 38.9 | 0 | 7 | 3 |
| Untreated 2 | 6.1 | **39.6** | 0 | 1 | 0 |
| Untreated 3 | 6.5 | 39.2 | 0 | 1 | 1 |
| Untreated 4 | 3.4 | 38.9 | 0 | 3 | 0 |
| Mupirocin treated 1 | 6.5 | 38.6 | 1 | 8 | 1 |
| Mupirocin treated 2 | 12.0 | **39.6** | 0.5 | 7 | 0 |
| Mupirocin treated 3 | 6.2 | **39.6** | 0 | 5 | 0 |
| Mupirocin treated 4 | 2.2 | 38.4 | 0 | 4 | 0 |
| Mupirocin treated 5 | 1.6 | 38.7 | 0 | 3 | 1 |
| Mupirocin treated 6 | 0.9 | **39.7** | 0 | 4 | 1 |
| Mupirocin treated 7 | 7.6 | 38.5 | 0 | 4 | 0 |
| Mupirocin treated 8 | 0.4 | 39.0 | 0 | 4 | 0 |
| *S.pneumoniae* infected 1 | 7 | 39.2 | 1 | 7 | 4 |
| *S.pneumoniae* infected 2 | 2.6 | 39.4 | 1 | 6 | 4 |
| *S.pneumoniae* infected 3 | 8.0 | **40.3** | 1 | 9 | 4 |
| *S.pneumoniae* infected 4 | 23.3 | **39.8** | 0 | 9 | 5 |
| *S.pneumoniae* infected 5 | 20.1 | 39.4 | 0 | 7 | 5 |

[a] Fever defined by temperature greater than 39.5°C
[b] Lethargy score: 0= alert and playful, 1=alert, playful when stimulated, 2=alert, not playful when stimulated, 3=neither alert nor playful when stimulated
